# Endogenous–Exogenous Competition Delays Internal Selection Across Cortical and Oculomotor Systems

**DOI:** 10.64898/2025.12.02.691938

**Authors:** Dayana Valdez Izquierdo, Edward F. Ester

## Abstract

Working memory (WM) enables the temporary maintenance of information for guiding thought and action, but its limited capacity requires internal selective attention mechanisms to prioritize goal-relevant representations. Like external attention, internal selection can be shaped by both endogenous (goal-driven) and exogenous (stimulus-driven) factors. Classic work by Charles Folk and Roger Remington demonstrated that such factors interact competitively during external attention, producing stimulus-driven capture under certain conditions. Whether similar competitive dynamics govern internal selection remains unclear. Prior studies using retro-cue paradigms with extended cue–response delays have reported evidence for “retro-capture”—involuntary orienting toward task-irrelevant but cue-matching working memory content. Here, we combined EEG and eye-tracking to test whether retro-capture persists when the cue and response probe are simultaneous, eliminating the temporal window in which capture has previously been observed. Participants remembered two oriented bars and used either a pro-cue or an anti-cue to guide recall. Across behavioral, cortical, and oculomotor measures, we found no evidence for retro-capture. Instead, competition between endogenous and exogenous selection mechanisms uniformly delayed the selection of task-relevant WM content, with delays evident in response times, lateralized alpha-band activity, and gaze biases toward remembered locations. Our findings indicate that retro-capture is not an obligatory consequence of endogenous–exogenous competition in WM but may depend on the temporal separation between cue and action.

---

Efficient behavior requires mechanisms for selecting information that is relevant to current goals while suppressing competing inputs. Foundational work by Charles Folk and Roger Remington—whose influential ideas this special issue celebrates—demonstrated that external attention is shaped by the interplay of endogenous (goal-driven) and exogenous (stimulus-driven) factors. Their contingent involuntary orienting framework showed that stimulus-driven capture depends critically on the match between stimulus properties and an observer’s attentional control settings, revealing that attentional competition unfolds through dynamic interactions between goals and salience rather than through purely bottom-up or top-down routes (Folk et al., 1992; Folk & Remington, 1998). This framework has profoundly shaped theories of attentional control and continues to inspire new lines of inquiry.

A central question in contemporary cognitive neuroscience is whether the same competitive principles govern selection within working memory (WM). Because WM is capacity-limited, internal attention must determine which representations gain privileged access to behavior and which are deprioritized or suppressed (Souza & Oberauer, 2016; Ester et al., 2018). Although internal attention shares functional similarities with external attention, it operates on stored rather than incoming information and may therefore rely on distinct neural mechanisms and computational constraints. Recent studies offer differing views on how endogenous and exogenous factors interact during internal selection. An eye-tracking study by van Ede et al. (2020) reported evidence for “retro-capture”: salient retrocues biased gaze toward matching but task-irrelevant WM content, suggesting that involuntary influences can briefly dominate internal selection. In contrast, an EEG study using a similar pro-cue/anti-cue manipulation (Ester & Nouri, 2023) found no evidence for retro-capture. Instead, competition between endogenous and exogenous selection mechanisms delayed the endogenous selection of task-relevant WM content without producing erroneous prioritization of irrelevant representations. Together, these findings leave unresolved whether retro-capture reflects a general property of endogenous–exogenous competition during internal selection or whether it depends on specific features of the task design.

Importantly, the studies that have examined endogenous–exogenous competition during internal selection share a common temporal structure: a retrocue is presented during the maintenance interval, followed by a blank delay of one to two seconds before any behavioral response is required. In van Ede et al. (2020), a colored retrocue appeared 1 second into a 3-second delay, leaving approximately 2 seconds before the probe. Ester and Nouri (2023) employed a similar mid-delay retrocue followed by a ∼1-second blank interval before the recall probe. This temporal separation between the cue and the response affords an extended window during which competitive selection dynamics—including involuntary shifts toward cue-matching but task-irrelevant content—can unfold prior to action. Note that in both studies, and in more recent work by Ding et al. (2024), the pro- and anti-cue conditions were blocked, meaning that participants maintained the task rule throughout a given block of trials rather than switching rules on a trial-by-trial basis (van Ede et al., 2020; Ding et al., 2024).

The present study departs from this common design in a theoretically important way. Here, the retrocue and probe are one and the same event: a color change of the central fixation cross at the end of the retention interval simultaneously informs participants which bar is task-relevant and signals them to begin their motor response immediately. There is no blank delay between the onset of the cue and the onset of recall. This design collapses the temporal separation between selection and action that characterized prior work, placing immediate motor demands on the selection process. Prior research has shown that when WM items are linked to specific actions, visual and motor selection occur concurrently and engage appropriate visual and motor brain areas in parallel (van Ede et al., 2019b). Furthermore, internal prioritization may proceed in stages—first orienting to a memorized representation, then reconfiguring it for upcoming task demands (Myers et al., 2017)—and immediate action requirements may compress or bypass the earlier orienting stage in which competitive dynamics manifest. Thus, our design provides a direct test of whether retro-capture—and competitive selection dynamics more broadly—depend on the temporal window afforded by extended cue-probe delays, or whether they reflect obligatory consequences of endogenous–exogenous competition that emerge regardless of action demands.

To address this question, we recorded EEG and eye-tracking data while participants performed a retrospectively cued visuomotor WM task. As in prior work (van Ede et al., 2020; Ester & Nouri, 2023; Ding et al., 2024), we manipulated whether retrocues matched task-relevant or task-irrelevant WM content, placing endogenous and exogenous selection mechanisms in alignment (pro-cue) or in competition (anti-cue). Several outcomes were possible. First, if retro-capture is an obligatory consequence of endogenous–exogenous competition in WM, it should emerge in both cortical and oculomotor signatures even when the cue and probe are simultaneous. Second, if retro-capture depends on the extended temporal window between cue and response that characterized prior designs, then eliminating this window should abolish capture while competition may instead manifest as delayed selection of task-relevant content. Third, even if retro-capture is absent, the magnitude of any selection delay may differ across cortical and oculomotor systems, which would suggest that these measures index functionally distinct stages of internal selection—with cortical oscillations reflecting the computation of attentional priority and gaze reflecting a downstream expression of the selected representation. By situating internal selection within the broader theoretical tradition shaped by Folk and Remington, our work asks not only whether endogenous and exogenous influences compete within WM, but also how the temporal demands of action shape the form that competition takes across physiological systems.

## Methods

### Data & Code Availability

Stimulus presentation software, EEG preprocessing routines, preprocessed EEG data, and analytic software sufficient to reproduce each manuscript figure and table are available via the Open Sciences Framework: https://doi.org/10.17605/OSF.IO/U8XN2

### Participants

35 human volunteers (both sexes) participated in this study. Volunteers aged 18-40 were recruited from the University of Nevada, Reno (UNR) community and completed a single 2.5-hour testing session in exchange for course credit or monetary compensation ($15/h). We had no a priori reason to expect that task performance would vary across sex, gender identity, race, ethnicity, handedness, or any other immutable characteristic. Thus, to better protect participants’ privacy, we did not collect this information. All experimental procedures were approved by the UNR Institutional Review Board. Data from 5 participants were discarded due to technical issues, for example, hardware failures that prevented event markers from being sent from the stimulus presentation computer to the EEG recording computer. The data reported here reflects the remaining 30 participants.

Our target sample size of 30 participants was selected to match or exceed those used in comparable studies examining endogenous–exogenous competition during internal selection (van Ede et al., 2020: N = 25; Ester & Nouri, 2023: N = 40; Ding et al., 2024: N = 25). A formal a priori power analysis was not performed because the relevant effect sizes for our novel task design (particularly the oculomotor measures under immediate response demands) were unknown at the time of data collection. We instead report a sensitivity analysis: with N = 30, our nonparametric bootstrap and permutation tests had 80% power to detect effects of Cohen’s d ≥ 0.53 (two-tailed, α = .05). For oculomotor analyses (N = 21), the corresponding sensitivity threshold was d ≥ 0.64. The reduced eye-tracking sample reflects participant-level factors (e.g., corrective lenses, mascara) that prevented acquisition of high-quality binocular gaze data and is acknowledged as a limitation.

### Testing Environment

Participants were seated in a dimly lit room for the duration of testing. Visual displays were generated in MATLAB and rendered on a 27-in. LCD monitor (1920 × 1080 resolution) cycling at 240 Hz using Psychtoolbox-3 (Kleiner, Brainard, & Pelli, 2007). Participants were seated 85 cm from the display via a chin rest and responded using a standard U.S. computer keyboard synced to the monitor refresh rate.

### Visuomotor Memory Tasks

Participants performed two variants of a visuomotor memory task used in prior research (van Ede et al., 2019; van Ede et al., 2020; Figure 1A). Each trial began with a 0.5 sec sample display containing colored bars (5.7° length x 0.8° height from a viewing distance of 85 cm) presented 5.7° to the left and right of a central fixation cross. The orientation of one bar was chosen from the interval -70°:1°:-20° (i.e., counterclockwise) and the orientation of the other bar was chosen from the interval 20°:1°:70° (i.e., clockwise). Thus, the sample display always contained one clockwise and one counterclockwise-tilted bar, though their positions were randomized over trials. The sample display was followed by a 1.25-1.75 sec blank delay interval, with the specific interval randomly and independently chosen from a uniform distribution with a 5 ms resolution on each trial. At the end of the trial, a probe, defined as a change in the color of the central fixation cross, instructed participants to recall the orientation of the matching bar. Importantly, the vertical tilt of the to-be-recalled bar determined which hand should be used for recall, with counterclockwise bars requiring a left-hand response and clockwise bars requiring a right-hand response. Participants recalled counterclockwise bars by pressing and holding the “z” key with their left index finger. Upon their initial response, a recall stimulus consisting of a circle with two endpoints appeared and began to rotate counterclockwise. Participants held down the z key until the orientation of the recall stimulus matched their memory for the orientation of the to-be-recalled bar. Likewise, participants recalled clockwise bars by pressing and holding the “?” key and releasing it when the recall stimulus matched their memory. We recorded three variables: accuracy, defined as the proportion of trials where the participant used the appropriate hand for recall, error, defined as the absolute angular difference between the reported and actual bar orientations, and response initiation time, defined as the interval between probe onset and participants’ initial button press response. During the pro-cue task, participants were instructed to recall the orientation of the bar matching the probe color. During the anti-cue task, participants were instructed to recall the orientation of the bar that did not match the probe color. Participants performed the pro- and anti-cue tasks sequentially, for example, 8 blocks of the pro-cue task followed by 8 blocks of the anti-cue task, though task order was counterbalanced across participants. Participants performed 4-10 (median 8) blocks of 48 trials in the pro-cue task and 6-10 blocks (median 8) of 48 trials in the anti-cue task.

**Figure 1.**
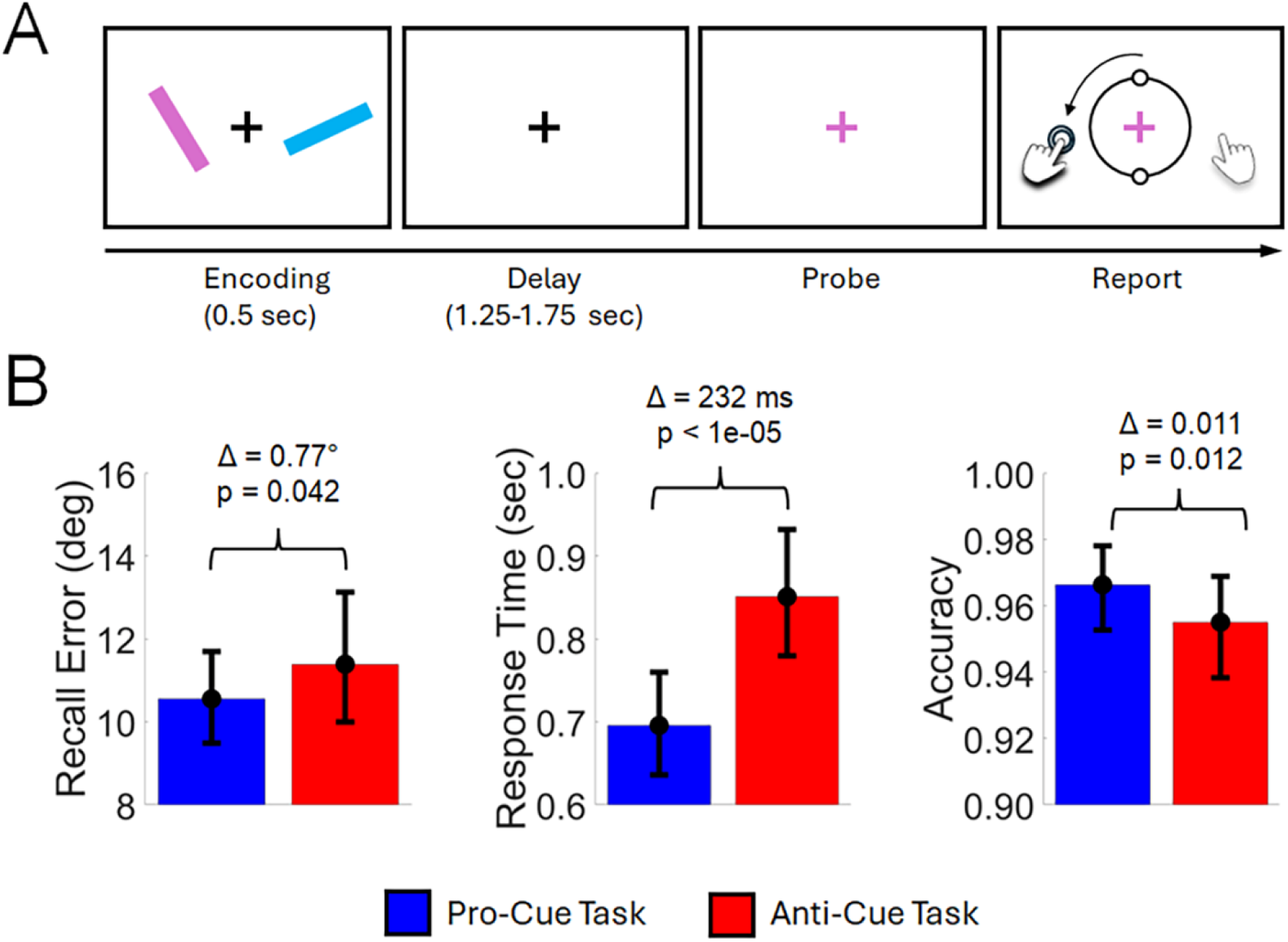
Visuomotor Task and Behavioral Performance. (A) Participants remembered two bars over a short delay, then recalled the orientation of a retrospectively probed bar by adjusting a report stimulus. During the pro-cue task, participants were tasked with reporting the orientation of the bar whose color matched the recall probe; during the anti-cue task, participants were tasked with reporting the orientation of the bar whose color did not match the recall probe. See Methods for additional details. (B) Recall error, response times, and accuracy plotted as a function of task. Accuracy refers to trials where the participant recalled the probed bar with the correct response hand.

### EEG Acquisition and Preprocessing

Continuous EEG was recorded from 63 scalp electrodes using a BrainProducts actiCHamp system. Online recordings were referenced to 10-20 site FCz and digitized at 1 kHz. The following offline preprocessing steps were applied, in order: (1) resampling from 1 kHz to 250 Hz, (2) high-pass filtering (0.5 Hz using zero-phase forward- and reverse-finite impulse response filters as implemented by EEGLAB software extensions; Delorme & Makeig, 2004), (3) identification and reconstruction of noisy electrodes and epochs via artifact subspace reconstruction (implemented via EEGLAB; Chang, Hsu, Pion-Tonachini, & Jung 2020), (4) re-referencing to the average of the left and right mastoids (operationalized as 10-20 sites TP9 and TP10), (5) epoching from −1.0 to +3.0 sec relative to probe onset, (6) interpolation of noisy electrodes, and (7) application of a surface Laplacian to remove low spatial frequency components from the signal (Perrin, Pernier, Bertrand, & Echallier, 1989).

### Eyetracking Acquisition and Preprocessing

We obtained high-quality binocular eye position data for 21 of 30 participants. It was not possible to acquire eye position data from the remaining 9 participants due to participant-level variables (e.g., participants who wore corrective lenses or mascara during their testing session). Eye position data were acquired using an SR Research Eyelink 1000 Plus infrared eye tracker and digitized at 500 Hz or 1 kHz. The eye tracker was calibrated at the beginning of each task (i.e., pro-cue and anti-cue) and drift correction was applied at the beginning of each task block. Data were resampled to 250 Hz and filtered for blinks. Blinks were defined as periods where the recorded pupil size fell below the median pupil size recorded across all experimental trials and conditions. We additionally discarded data from 50 ms before and 50 ms after each blink. Missing data were linearly interpolated, and the resulting gaze timeseries were smoothed with a gaussian kernel (10 ms full-width half-maximum). No other preprocessing steps were applied.

### Spectral EEG Analyses

Spectral analyses focused on occipitoparietal 10–20 electrode site pairs O1/2, PO3/4, PO7/8. These sites were selected based on prior work demonstrating that lateralized alpha-band modulations during spatial attention and working memory selection are maximal over posterior occipitoparietal scalp regions (Gould et al., 2011; Bacigalupo & Luck, 2019; Ester & Weese, 2023). Analyses were restricted to a period spanning −0.5 to 1.5 around probe onset. We extracted broadband spectral power from each electrode on each trial using a short-time Fourier transform (STFT) with a frequency range of 1–40 Hz (in 0.25-Hz steps) using a 200-msec sliding window and 10-msec step size. Power was computed by squaring the absolute value of the complex Fourier coefficients within each STFT window. To quantify changes in lateralized activity during visual selection, we sorted power estimates at each occipitoparietal electrode site by the location of the retrospectively cued stimulus, that is, contralateral versus ipsilateral hemifield, and expressed this difference as a normalized percentage:

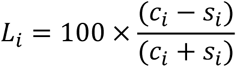

where *ci* and *si* are the trial-averaged responses of electrode *i* when the probe stimulus appeared in the contralateral or ipsilateral visual field, respectively. Lateralization estimates were pooled across occipitoparietal electrode sites, yielding a single time frequency matrix of power estimates per participant.

To generate topographical maps of lateralized power relative to the probed stimulus location (i.e., left vs. right visual hemifield), we repeated this analysis for every scalp electrode and averaged power estimates over 8–13 Hz (“alpha-band” activity) during a period spanning of 500–1000 msec following probe onset. This frequency band was chosen a priori based on commonly reported values in the literature (e.g., Klimesch, 2012), and this specific temporal window was chosen a priori based on prior reports suggesting that it takes human observers 300–500 msec to process and respond to a retrospective memory cue (e.g., Souza et al., 2014). To visualize changes in occipitoparietal alpha power over time we extracted and averaged lateralization estimates from 8 to 13 Hz over electrode sites O1/2, PO3/4, and PO7/8, Statistical analyses of lateralization were performed using nonparametric sign permutation tests with temporal cluster-based corrections (see Statistical Comparisons section).

### Gaze Bias Analyses

Preprocessed eye position data were sorted by the location of the retrospectively cued stimulus (i.e., left vs. right visual field) and task (i.e., pro- vs. anti-cue). Because stimuli were rendered along the vertical meridian, we restricted our analyses to horizontal eye position measurements. Following earlier work (van Ede et al. 2019), we computed a normalized measure of gaze bias by converting pixelwise recordings into a percentage along an axis extending from fixation to the center of each visual stimulus. Thus, a 100% gaze bias would result from participants foveating the center of one stimulus (with negative and positive values corresponding to gaze shifts towards the left and right visual hemifields, respectively), whereas a 0% gaze bias would result from participants perfectly holding fixation.

### Statistical Comparisons – Behavior

Three features of our dependent variables motivated the use of nonparametric bootstrap tests rather than standard parametric alternatives (e.g., t-tests). First, response time distributions are known to be positively skewed, typically following an ex-Gaussian rather than a Gaussian form (Luce, 1986; Whelan, 2008). Second, recall errors were expected to cluster near the lower bound of the scale (i.e., 0°), and accuracy rates near the upper bound (i.e., 100%), producing floor and ceiling effects that compress variability and violate assumptions of normality (Šimkovic & Träuble, 2019). Because the sample mean is a biased estimator of central tendency for skewed distributions with small-to-moderate sample sizes (Miller, 1988), and because parametric tests assume normally distributed sampling distributions of the test statistic, we adopted bootstrap resampling procedures that make no distributional assumptions (Efron & Tibshirani, 1993). For each dependent variable (e.g., recall error) and experimental condition (e.g., pro-cue task) we randomly selected, with replacement, and averaged data from 30 of 30 volunteers. This procedure was repeated 10,000 times, yielding 10,000-element distributions of sample means. Next, we compared the number of permutations where the estimated mean in one condition (e.g., the pro-cue task) exceeded the mean in the other condition (e.g., the anti-cue task), yielding an empirical p-value.

### Statistical Comparisons – EEG and Eyetracking

Statistical comparisons were based on nonparametric signed randomization tests (Maris & Oostenveld, 2007). Unless otherwise specified, each test we performed assumes a null statistic of 0 (i.e., no difference in alpha-band lateralization or no detectable gaze bias). We generated null distributions by randomly relabeling each participant’s data with 50% probability and averaging the data across participants. This step was repeated 10,000 times, yielding a 10,000-element null distribution for each time point. Finally, we implemented a cluster-based permutation test with cluster-forming and cluster size thresholds of p < .05 (two-tailed) to evaluate observed differences with respect to the null distribution while accounting for signal autocorrelation.

Task-dependent differences in the latency of visual selection were evaluated with jackknife tests (Miller, Patterson, & Ulrich, 1998; Kiesel, Miller, Jolicoeur, & Brisson, 2008). For EEG and gaze position data, we selected and averaged data from N – 1 participants, then computed the time at which the absolute value of lateralization was greatest. This procedure repeated until every participant’s data has been held out, yielding a 30-element (EEG) or 21-element (Gaze Bias) vector of latencies for each task (i.e., pro- vs. anti-cue). For each jackknife sub-average, we computed three latency metrics: onset latency (the first time point at which the waveform reached 50% of its peak amplitude), 50% area latency (the time point dividing the total signed area under the lateralization waveform into equal halves), and peak latency (the time of maximum negative lateralization). We compared latency estimates across tasks via non-parametric bootstrap tests. We created distributions of pro- and anti-cue latencies by randomly selecting (with replacement) and averaging latency estimates 10,000 times. We subtracted the distribution of pro-cue latencies from the anti-cue latencies and calculated the number of times where latency estimates for the anti-cue condition were less than those for the pro-cue condition. This approach yields an empirical p-value with a minimum of 1e-04.

### Brain–Behavior Correlations

To evaluate whether individual differences in the behavioral cost of competition (response time slowing) were related to individual differences in the cortical selection delay (alpha lateralization latency), we correlated the across-condition difference in response times (anti-cue minus pro-cue) with the across-condition difference in alpha lateralization latency. Individual-participant latency estimates were recovered from jackknife sub-averages using the bias-corrected transform described by Smulders (2010): 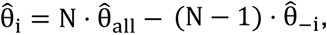 where 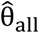 is the latency estimated from the full group-average waveform, 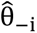 is the latency estimated from the sub-average excluding participant i, and N is the sample size. This procedure was applied separately for each of three latency metrics: onset latency (first time point reaching 50% of peak amplitude), 50% area latency (time point dividing the total negative area into equal halves), and peak latency (time of maximum negative lateralization). Correlations were computed using Spearman’s rank correlation to accommodate potential non-normality in the recovered latency estimates (Smulders, 2010).

### Simulation-Based Sensitivity Analysis

To evaluate whether retro-capture might occur on a subset of anti-cue trials but be obscured by trial averaging, we performed a simulation-based sensitivity analysis. We constructed two template waveforms: a “delayed selection” template derived from the observed group-average anti-cue alpha lateralization, and a “retro-capture” template consisting of the delayed-selection waveform plus a transient positive deflection (toward the task-irrelevant, cue-matching item) during the first 500 ms post-probe. The amplitude of this transient was scaled as a proportion of the observed pro-cue peak lateralization (10%, 15%, or 20%), consistent with the relative magnitudes of involuntary and voluntary gaze biases observed in prior retro-cueing studies (van Ede et al., 2019a; Ding, Postle, & van Ede, 2024). For each simulation, we generated synthetic participant-averaged anti-cue waveforms for 30 participants, where a specified proportion of trials (0–100%, in 5% increments) followed the retro-capture template and the remaining trials followed the delayed-selection template. Participant-level noise was estimated from the between-subject residuals of the observed anti-cue data, preserving the temporal structure of the noise. We then tested whether the early post-probe window (0–500 ms) showed a significant positive deflection relative to the delayed-selection baseline using a one-tailed one-sample t-test. This procedure was repeated 1,000 times per mixture proportion to estimate statistical power as a function of retro-capture prevalence.

We performed an analogous simulation for the gaze bias data. We constructed a delayed-selection gaze template from the observed group-average anti-cue gaze lateralization and a retro-capture template consisting of the delayed-selection waveform plus a transient negative deflection (toward the task-irrelevant item) during the first 500 ms post-probe, scaled as 10%, 15%, or 20% of the observed pro-cue peak gaze bias. Synthetic participant-averaged waveforms were generated for 21 participants, with participant-level noise estimated from the between-subject residuals of the observed anti-cue gaze data. Detection was assessed via a one-tailed one-sample t-test.

## Results

### Overview

We recorded EEG and horizontal gaze position data while human volunteers performed a retrospectively cued visuomotor working memory task (Figure 1A). Each trial began with a 0.5-sec sample display containing two colored bars (one clockwise-tilted, one counterclockwise-tilted) presented to the left and right of central fixation. Following a 1.25–1.75 sec blank delay, a probe stimulus (a color change of the central fixation cross) instructed participants which bar to recall. Critically, the vertical tilt of the to-be-recalled bar determined which hand should be used for recall: counterclockwise bars required left-hand responses, whereas clockwise bars required right-hand responses. During the pro-cue task, participants recalled the orientation of the bar matching the probe color. During the anti-cue task, participants recalled the orientation of the bar that did not match the probe color. Thus, in pro-cue blocks, endogenous (goal-driven) and exogenous (stimulus-driven) selection factors were aligned, whereas in anti-cue blocks these factors competed.

We evaluated two models, which we term retro-capture and delayed-selection. The retro-capture account predicts that competition between endogenous and exogenous selection mechanisms elicit an involuntary shift of attention towards the location of the cue-matching but task-irrelevant bar during the anti-cue task, and thus, both EEG and eye-tracking measurements of internal selection should exhibit an initial bias towards the location of the cue-matching but task-irrelevant bar during the anti-cue task. The delayed selection account predicts that competition between endogenous and exogenous selection mechanisms does not elicit an involuntary shift of attention but engages attention control mechanisms to resolve this competition, delaying the selection of task-relevant WM content. Thus, EEG and eye-tracking measurements of internal selection should be delayed during the anti-cue compared to the pro-cue task but should not exhibit a bias towards the task-irrelevant stimulus.

To test these alternatives, we defined EEG and eye-tracking measures of internal selection time-locked to the onset of the recall probe as follows: For EEG, extracted and calculated alpha power (8-13 Hz) over occipitoparietal electrode sites contralateral and ipsilateral to the visual hemifield containing the task-relevant bar. Based on prior work (Klimesch et al., 2012; van Ede et al., 2019a) we defined selection as the normalized difference in alpha power over electrode sites contralateral and ipsilateral to the location of the task-relevant stimulus (see Methods), where negative values reflect shifts of attention towards the task-relevant stimulus while positive values are assumed to reflect shifts of attention towards the task-irrelevant stimulus. For eye tracking, we extracted and converted records of participants’ horizontal gaze position into a normalized metric where 0% represents fixation and 100% represents foveating the task-relevant bar (i.e., cue-matching during the pro-cue task, and cue-nonmatching in the anti-cue task.

### Behavioral Performance

We quantified task performance using three measures: recall error (mean absolute angular difference between reported and actual bar orientations), response time (interval between probe onset and initial button press), and accuracy (proportion of trials using the correct response hand). Behavioral data are shown in Figures 1B–D. Recall error was modestly but significantly lower during the pro-cue task compared to the anti-cue task (M = 11.77° vs. 12.54°; p = 0.042; Figure 1B). Likewise, accuracy was slightly higher during the pro-cue versus anti-cue task (M = 96.63% vs. 95.52%; p = 0.012; Figure 1B). Most notably, response times were substantially slower during the anti-cue task compared to the pro-cue task (M = 0.962 sec vs. 0.730 sec; Δ = 232 ms; p < 1e-04; Figure 1B). This robust latency effect replicates prior findings (Ester & Nouri, 2023) and confirms that placing endogenous and exogenous selection slows participants’ responses.

### Alpha-band EEG Signatures of Internal Selection

Figure 2A summarizes lateralized changes in spectral power (i.e., contralateral to the task-relevant stimulus minus ipsilateral to the task-relevant stimulus) time-locked to probe onset and pooled over the pro- and anti-cue tasks. We identified a significant cluster of lateralized activity spanning ∼8-13 Hz and ∼0.5 to 1.0 sec after probe onset. This activity was clustered over occipitoparietal electrode sites, replicating prior findings (van Ede et al., 2019; Ester et al., 2023; Figure 2A, inset). Figure 2B plots lateralized alpha power from 8-13 Hz calculated from posterior electrode sites as a function of task. We found no evidence for selection of the cue-matching but task-irrelevant stimulus during the anti-cue task (i.e., positive lateralization values). Instead, we observed clear evidence for selection of the task-relevant stimulus, with the onset of selection significantly faster during the pro-cue task (M = 386 ms) vs. the anti-cue task (M = 503 ms; p < 1e-04; jackknife test). This pattern held when we defined latency as the 50% fractional area latency (674 vs. 862 ms; p < 1e-04) and peak negative alpha lateralization (609 vs. 763 ms; p < 1e-04; Table 1). This pattern replicates findings by Ester and Nouri (2023) and indicates that competition between endogenous and exogenous selection mechanisms engages attention control mechanisms that delay the selection of task-relevant WM content.

**Figure 2.**
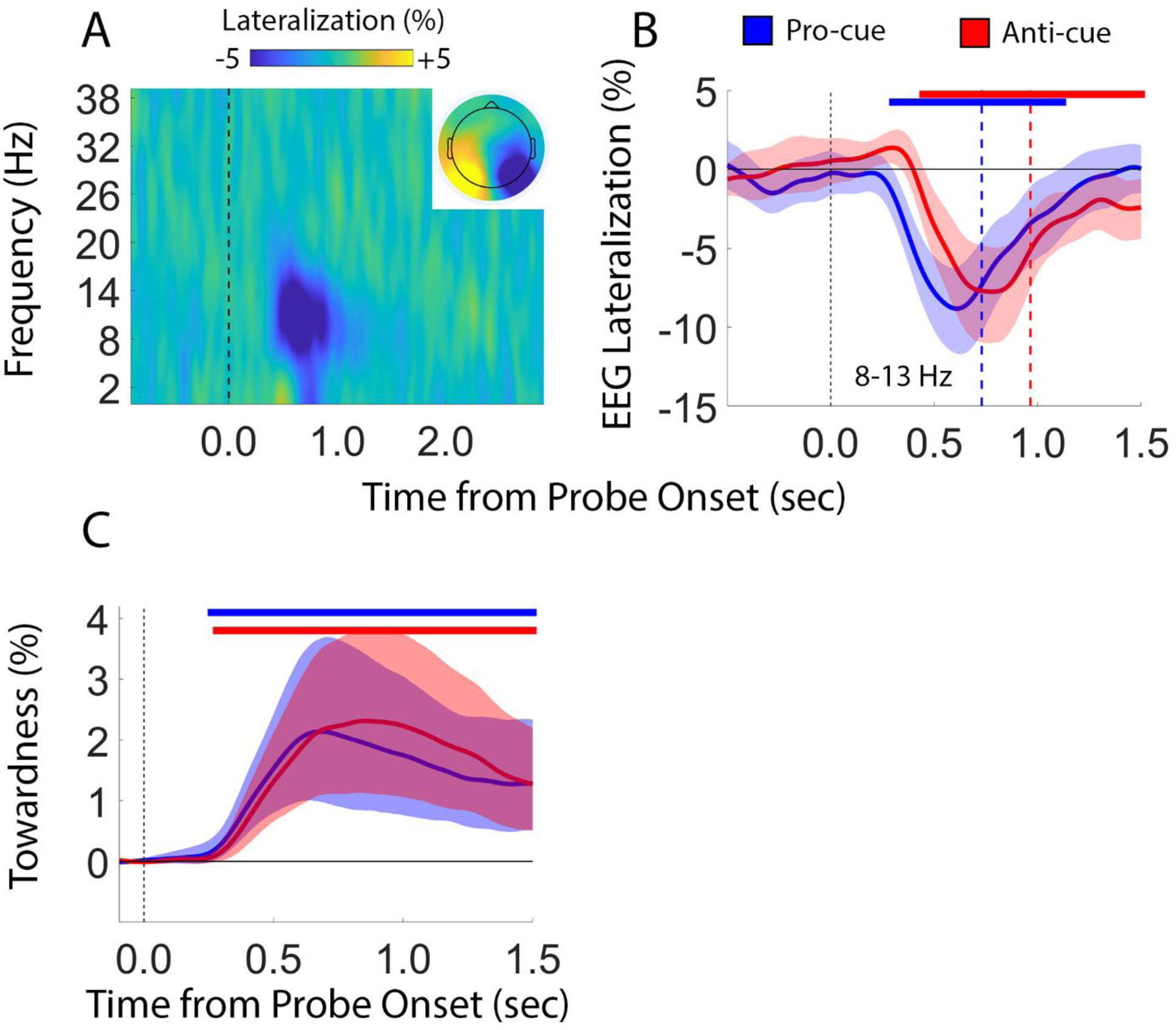
EEG and Oculomotor Signatures of Visual Selection. (A) Spectrogram plotting lateralized changes EEG activity as a function time-locked to probe onset, pooled across the pro- and anti-cue tasks. Inset shows the scalp distribution lateralized alpha band activity (8-13 Hz) over a period spanning 500-1000 ms after probe onset. (B) Lateralized alpha-band activity plotted as a function of task. (C) Lateralized gaze position plotted as a function of probe stimulus location (left vs. right visual field) and task (pro- vs. anti-cue). Shaded regions in panels B and C depict the 95% confidence interval of the mean. Solid and dashed bars at the top and bottom of panels B and C depict epochs where EEG and gaze lateralization were significantly different from 0. The vertical blue and red lines in panel B depict the mean response times during the pro- and anti-cue tasks, respectively.

**Table 1.** Alpha-lateralization Onset Estimates. AUC, 50% area latency: the time point at which 50% of the total signed area under the lateralization waveform has accumulated.

|  | Pro-Cue | Anti-Cue | Difference | p-value |
| --- | --- | --- | --- | --- |
| Onset Latency | 386 | 503 | 117 | < 1e-04 |
| 50% AUC | 674 | 862 | 188 | < 1e-04 |
| Peak Latency | 609 | 763 | 154 | < 1e-04 |

### Brain–Behavior Correlations

To test whether individual differences in the cortical selection delay predicted individual differences in behavioral slowing, we correlated the across-condition difference in response times (anti-cue minus pro-cue) with the across-condition difference in alpha lateralization latency for each of three latency metrics (see Methods). None of these correlations reached significance (onset latency: ρ = 0.184, p = 0.239; 50% area latency: ρ = 0.013, p = 0.946; peak latency: ρ = −0.222, p = 0.329; Table 2). Thus, while both behavioral and cortical measures showed robust group-level effects of competition, the magnitude of these effects was not correlated across participants. We return to this point in the discussion.

**Table 2.** Correlations Between Task-Level Differences in Alpha Lateralization Latency and Response Time Differences.

| | $\rho$ | p-value |
| --- | --- | --- |
| Onset Latency | 0.184 | 0.329 |
| 50% AUC | 0.013 | 0.946 |
| Peak Latency | -0.222 | 0.239 |

### Oculomotor Signatures of Internal Selection

Several recent studies have linked small (< 1°) biases in gaze position with the selection of information held in WM (e.g., van Ede et al., 2019a; Ester & Weese, 2023; Chota et al., 2025). In the context of the visuomotor WM task used here, the typical finding is that participants’ gaze is subtly but robustly biased toward the location of a task-relevant stimulus following an informative retrocue (e.g., van Ede et al., 2019a). Notably, van Ede et al. (2020) exploited these biases to test how competition between endogenous and exogenous factors influences the selection of task-relevant WM content. During an anti-cue task, they reported that participants’ gaze initially shifted toward the location of a cue-matching but task-irrelevant stimulus followed by a corrective gaze shift toward the task-relevant stimulus, an effect they termed “retro-capture.” Although we found no evidence for a retro-capture effect using EEG signatures of internal selection (Figures 2A–B), there is growing evidence that oculomotor and electrophysiological signatures of external and internal attention intersect in complex ways (e.g., Liu et al., 2022; Yu et al., 2022; Liu et al., 2025). Thus, we sought to test whether oculomotor retro-capture would emerge even in the absence of cortical retro-capture.

We found no evidence for oculomotor retro-capture: gaze did not shift toward the cue-matching but task-irrelevant stimulus in the anti-cue condition (Figure 2C). Instead, as with the EEG data, gaze biases toward the task-relevant stimulus were modestly but significantly delayed during the anti-cue compared to the pro-cue task (Table 3). However, the competition-induced delays in gaze onset were numerically smaller than the corresponding EEG delays (gaze: Δ = 51 ms onset, Δ = 58 ms 50% area latency; EEG: Δ = 117 ms onset, Δ = 188 ms 50% area latency), suggesting that oculomotor selection may be less sensitive to the initial time cost of endogenous–exogenous competition than cortical selection. Similarly, across tasks, gaze bias onset lagged alpha lateralization onset by approximately 70–140 ms, consistent with the idea that gaze reflects a downstream expression of the cortical selection signal.

**Table 3.** Gaze Lateralization Onset Estimates. AUC, 50% area latency: the time point at which 50% of the total signed area under the lateralization waveform has accumulated.

|  | Pro-Cue | Anti-Cue | Difference | p-value |
| --- | --- | --- | --- | --- |
| Onset Latency | 521 | 572 | 51 | < 1e-04 |
| 50% AUC | 966 | 1024 | 58 | < 1e-04 |
| Peak Latency | 777 | 955 | 178 | < 1e-04 |

### Control Analyses

We considered the possibility that retro-capture effects were obscured by pooling data across trials. To evaluate this possibility, we re-computed EEG lateralization (Figure 3) and gaze bias lateralization (Figure 4) after applying a median split to participants’ response times and recall errors during the pro- and anti-cue tasks. Neither measure was influenced by response times nor recall error, suggesting that trial averaging did not obscure critical effects.

**Figure 3.**
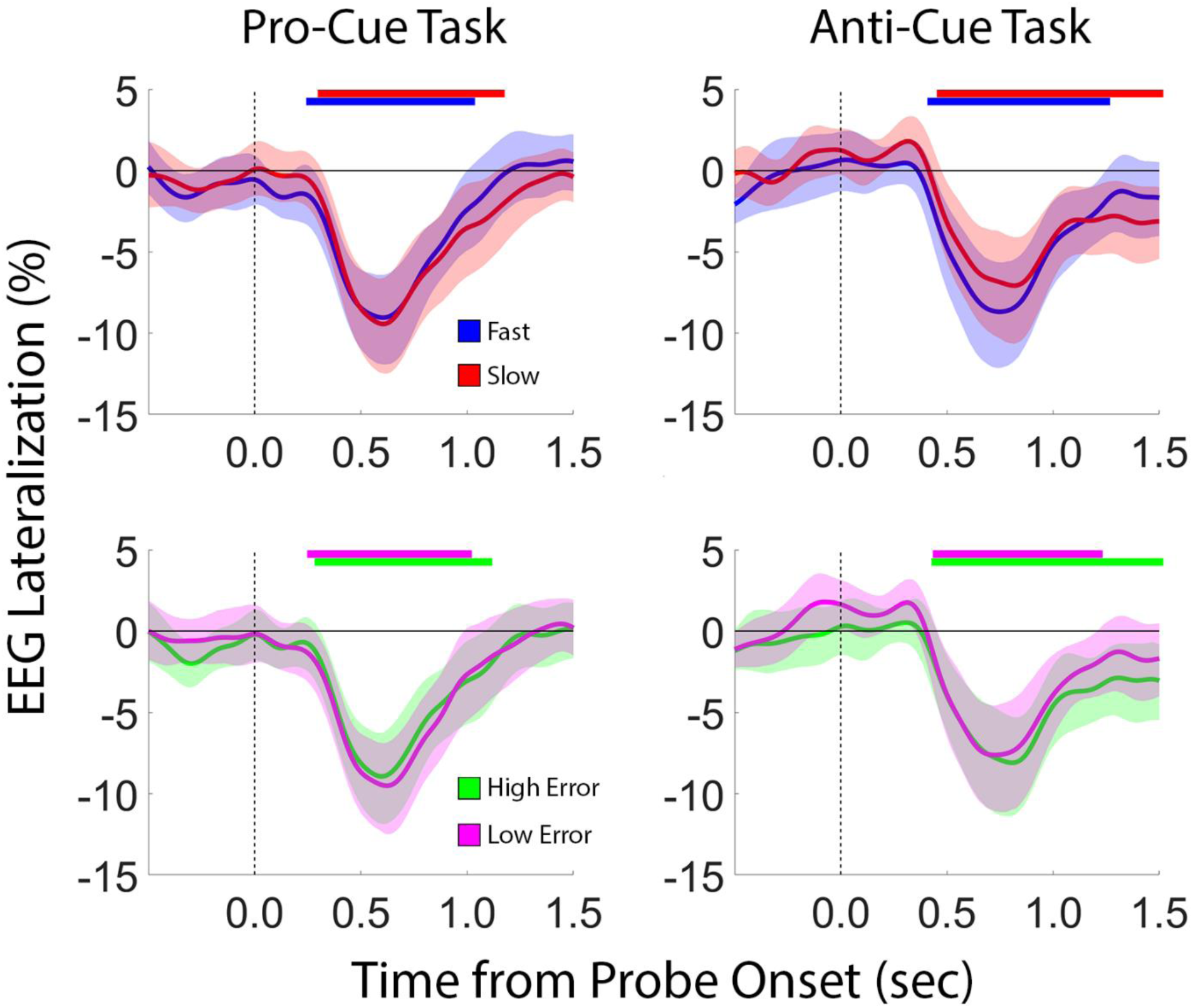
Lateralized EEG Activity Sorted by Task Performance. To test whether retro-capture effects were obscured by trial averaging, we recomputed EEG Lateralization during the pro-cue (left column) and anti-cue tasks (right column) after applying a median split to participants’ response times (top row) and recall error (bottom row). We found no evidence for retro-capture (i.e., lateralization values significantly greater than 0%).

**Figure 4.**
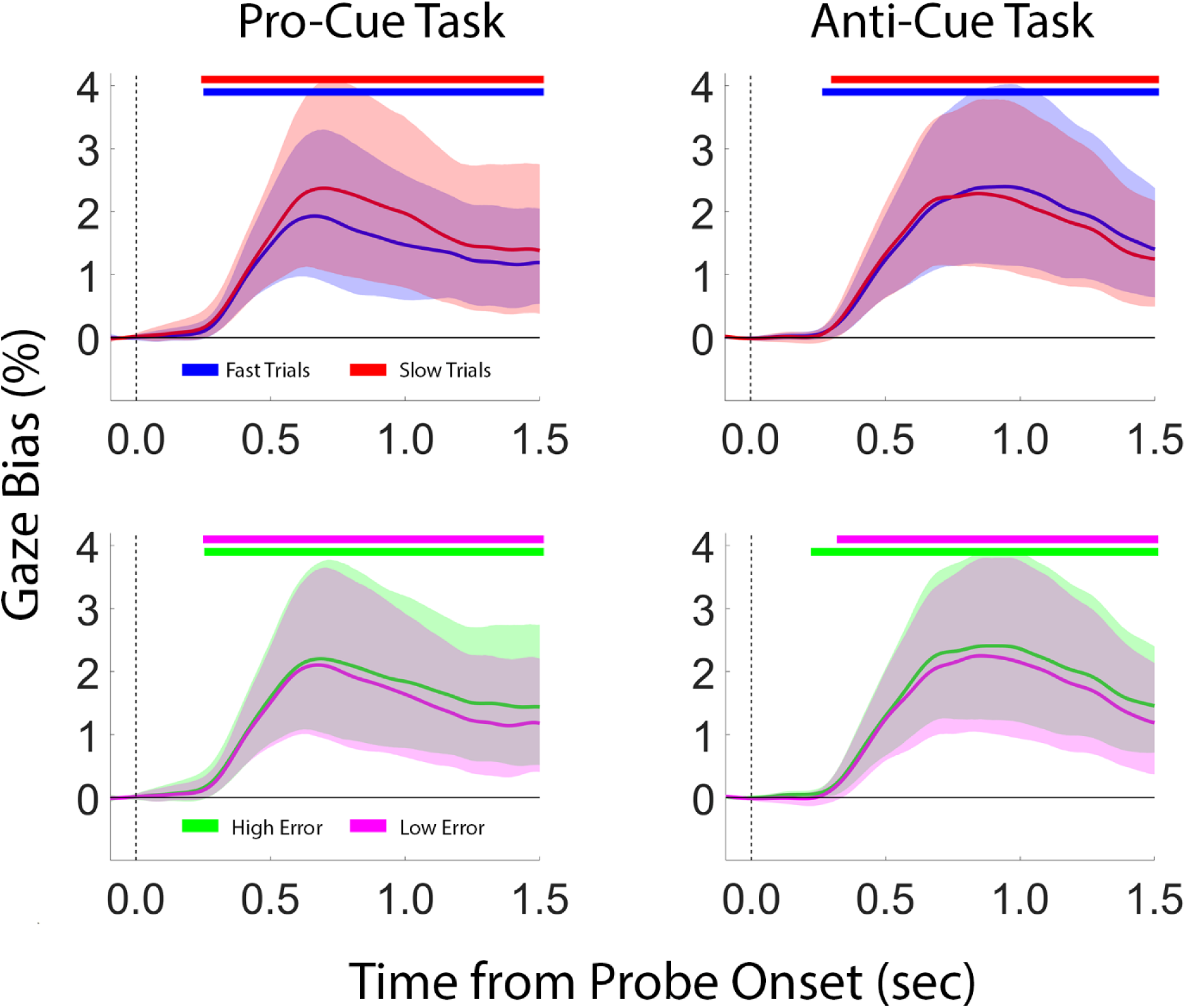
Gaze Bias Sorted by Task Performance. To test whether retro-capture effects were obscured by trial averaging, we recomputed gaze lateralization during the pro-cue (left column) and anti-cue tasks (right column) after applying a median split to participants’ response times (top row) and recall error (bottom row). We found no evidence for retro-capture (i.e., lateralization values significantly less than 0%).

Additionally, to evaluate whether retro-capture might occur on a subset of anti-cue trials but be obscured by trial averaging, we performed a simulation-based sensitivity analysis (see Methods). We constructed synthetic anti-cue alpha lateralization waveforms in which a specified proportion of trials (0–100%, in 5% increments) followed a “retro-capture” template—the observed delayed-selection waveform plus a transient positive deflection toward the task-irrelevant item during the first 500 ms post-probe—while the remaining trials followed the observed delayed-selection template. We tested the detectability of the early positive deflection across three assumed retro-capture amplitudes (10%, 15%, and 20% of the observed pro-cue peak lateralization), scaled to match the relative magnitudes of involuntary and voluntary gaze biases in prior work (van Ede et al., 2019a; Ding et al., 2024). This analysis revealed sensitivity to detect retro-capture effects on ∼20% of trials with 80% probability (Figure 5A). Thus, while we cannot exclude the possibility that retro-capture effects were obscured by trial averaging, we note that these effects would have to be very rare to produce the data reported here.

**Figure 5.**
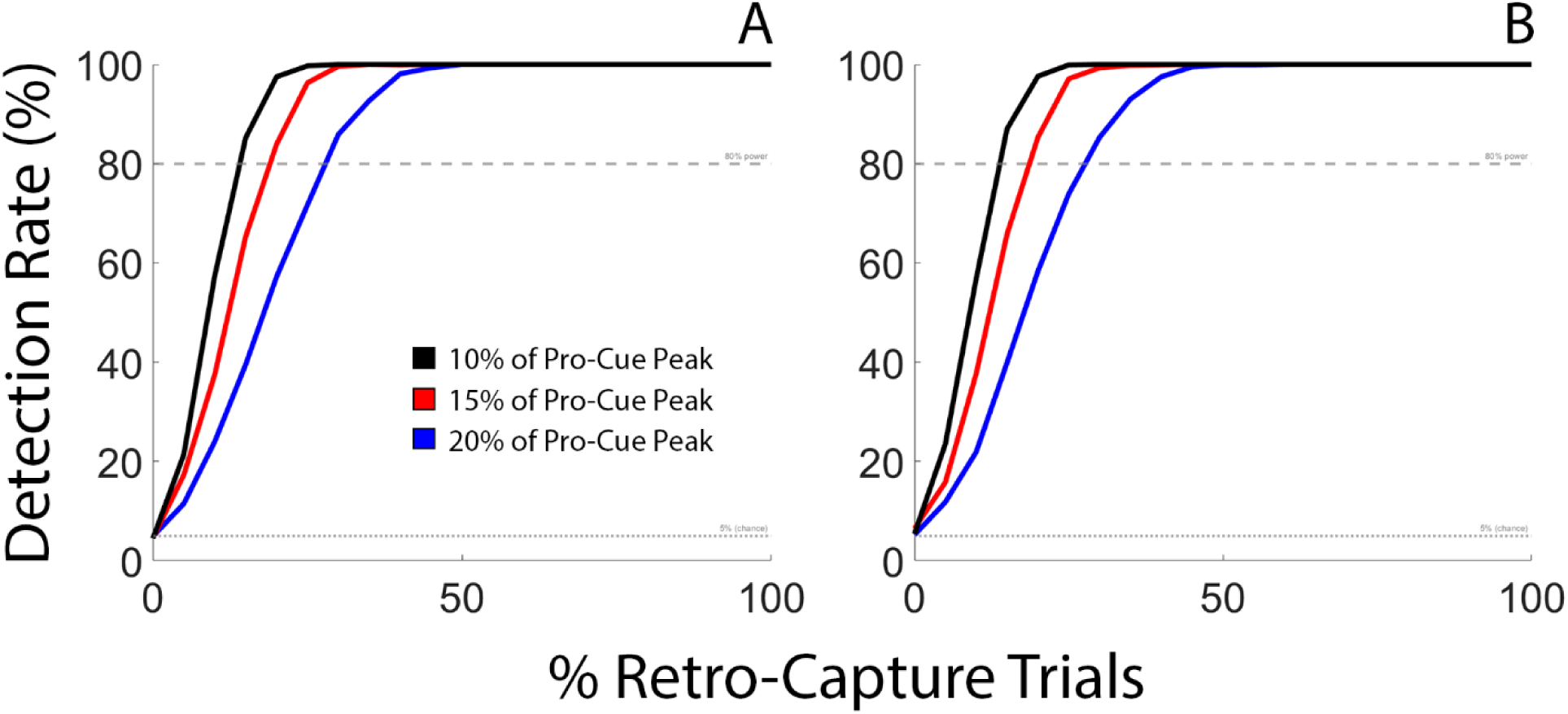
Simulation-based sensitivity analysis for retro-capture detection. (A) The probability of detecting a significant early positive deflection (toward the task-irrelevant item) in the trial-averaged alpha lateralization waveform is plotted as a function of the proportion of retro-capture trials, separately for three assumed capture amplitudes (10%, 15%, and 20% of the observed pro-cue peak lateralization). Horizontal dashed lines indicate 80% power and the 5% false positive rate. At empirically plausible capture amplitudes (10-15%), retro-capture would need to occur on ∼22-31% of anti-cue trials to be readily detected. (B) An analogous simulation was used for gaze position data.

A parallel simulation for gaze bias data yielded similar conclusions. At empirically plausible capture amplitudes (10–15% of the observed pro-cue peak gaze bias), retro-capture would need to occur on approximately 12–25% of anti-cue trials to be detected with 80% power, indicating adequate sensitivity to detect oculomotor retro-capture had it been present (Figure 5B).

## Discussion

The present study investigated how endogenous and exogenous influences jointly shape internal selection in WM, and whether competitive dynamics previously observed under extended cue–response delays generalize to conditions requiring immediate action. By combining EEG and eye-tracking during a visuomotor WM task in which the retrocue and response probe were simultaneous, we found convergent evidence that endogenous–exogenous competition delays the selection of task-relevant WM content without producing retro-capture. Delays were evident across behavioral responses, lateralized alpha-band activity, and oculomotor biases, though the magnitude of the delay was greater in cortical than oculomotor signatures. These findings indicate that retro-capture is not an obligatory consequence of endogenous–exogenous competition in WM and suggest that cortical and oculomotor systems, while both sensitive to competition, differ in the degree to which competitive dynamics influence their selection time course.

Alpha-band lateralization exhibited the largest and most robust delays during the anti-cue condition—consistent with prior evidence that competition prolongs the selection of goal-relevant content (Ester & Nouri, 2023). Oculomotor biases toward the task-relevant item showed a qualitatively similar but smaller delay, with no evidence for retro-capture. Consistent with this view, gaze bias onset reliably followed alpha lateralization onset by on the order of 100 ms, indicating that oculomotor selection inherits, rather than initiates, the competition-induced delay. This pattern is consistent with a hierarchical account in which cortical oscillations reflect the primary computation of attentional priority—and are therefore most sensitive to the time cost of resolving competition—while oculomotor outputs reflect a downstream expression of the selected representation that inherits the delay in attenuated form (Awh, Armstrong, & Moore, 2006; though we note that direct quantitative comparisons of latency estimates across EEG and gaze should be interpreted cautiously, given differences in signal type, temporal resolution, and sample size). By the time oculomotor planning systems receive input regarding which WM item is task-relevant, competition may have already been largely resolved at earlier cortical stages. This interpretation aligns with recent work indicating that microsaccade biases correlate with, but are not necessary for, oscillatory modulations during internal attention (Liu et al., 2022; Liu et al., 2025).

The complete absence of retro-capture contrasts with van Ede et al. (2020) and Ding et al. (2024), who reported oculomotor retro-capture in anti-cue paradigms with extended cue–response delays. In both prior studies, the retrocue appeared during the maintenance interval and was followed by a 1–2 second blank delay before the response, affording a temporal window in which involuntary orienting could unfold without immediate motor consequences. In the present study, the retrocue and response probe were simultaneous, requiring participants to respond immediately. This collapse of the cue–response interval may prevent retro-capture from manifesting, either because immediate motor demands engage top-down control that suppresses involuntary orienting, or because retro-capture requires a quiescent post-cue period to develop through passive biased-competition dynamics. These accounts make distinguishable predictions: if motor demands actively suppress capture, retro-capture should be absent whenever the cue signals an immediate response; if capture requires a quiescent period, reintroducing even a brief blank delay should restore it. Critically, although retro-capture was absent, delayed selection persisted—replicating and extending Ester and Nouri (2023) and demonstrating that the time cost of resolving competition is an intrinsic feature of internal selection that cannot be bypassed by immediate action demands. One interpretation is that retro-capture reflects a slow, passive process requiring time to develop, whereas delayed selection reflects the active computational cost of overriding a prepotent cue-driven signal regardless of when the response is required.

Although behavioral, cortical, and oculomotor measures all showed robust group-level effects of competition, we found no significant correlations between the across-condition difference in alpha lateralization latency and the corresponding difference in response times (Table 2). This pattern may partly reflect the inherent noisiness of per-participant latency estimates recovered from jackknife sub-averages, but it is also consistent with the view that response times reflect multiple processing stages beyond spatial selection—including rule application, motor preparation, and execution—that vary independently across participants. Recent work supports this interpretation: the magnitude of cue-induced alpha lateralization is uncorrelated with memory precision, guessing, and distractor bias in retro-cue tasks, suggesting that lateralized alpha indexes an automatic attentional shift that coincides with, but is not causally responsible for, mnemonic prioritization (Mössing & Busch, 2020; Balestrieri et al., 2024). Under this framework, the group-level alpha delay reflects a genuine effect of competition on spatial prioritization, whereas the absence of individual-differences correlations reflects additional processing stages interposed between selection and behavior.

A major contribution of this work is clarifying how internal selection resolves competition differently from external attention. In the external domain, competition often produces involuntary capture by salient but irrelevant stimuli (Folk & Remington, 1998; Theeuwes, 1992). Internal selection, by contrast, resolves competition through slower, controlled processes without errant prioritization of irrelevant content. One interpretation is that internal attention operates over already-encoded representations that are inherently more stable and less susceptible to fleeting salience signals, tolerating slower resolution processes that emphasize stability over speed. This view resonates with the Internal Dominance Over External Attention (IDEA) hypothesis (Verschooren & Egner, 2023), which posits an inherent bias toward internal information. Our findings suggest that such dominance may come at a computational cost: internal attention preserves goal-relevant content but resolves conflict more slowly. Rather than a single unified selection mechanism, our data support frameworks positing multiple, partially independent selection processes that operate at distinct stages—from cortical prioritization to oculomotor implementation—with the magnitude of competitive influence decreasing at later stages.

Several limitations temper these conclusions. First, usable eye-tracking data were obtained from only 21 of 30 participants, reducing power to detect subtle oculomotor effects. Second, our minimal gaze preprocessing pipeline may have obscured microsaccade-level modulations. Third, because our design merged the retrocue and probe, we cannot determine whether competitive oculomotor dynamics would emerge in this sample under a temporally extended design. Fourth, the blocked design means we cannot rule out the possibility that trial-by-trial rule switching in a mixed design would produce different competitive dynamics. Future work parametrically varying the cue-to-response interval within participants would directly test whether the temporal window between selection and action modulates competitive dynamics in oculomotor signatures.

Together, our findings highlight that internal selection is neither a simple extension nor a direct analogue of external attention, but is supported by multiple physiological systems that are jointly sensitive to endogenous–exogenous competition yet resolve it at different timescales. This work is deeply informed by the intellectual legacy of Drs. Charles Folk and Roger Remington, whose pioneering studies revealed how endogenous and exogenous influences jointly shape external selection. By extending this logic to the internal realm of working memory, our findings illustrate how their theoretical framework continues to inspire new questions. In showing that internal selection relies on hierarchically organized mechanisms that are differentially sensitive to competition, we hope to honor their enduring influence and contribute to the ongoing evolution of ideas they helped bring to the field.

## Notes

https://doi.org/10.17605/OSF.IO/U8XN2

### Competing Interest Statement

The authors have declared no competing interest.

### Summary of Updates

Added Figure 5 × Corresponding Analyses, revisions to text.

### Funding

- NSF: 2050833

